# *Trichoderma afroharzianum* behaves differently in interaction with pea plants under varying iron availability

**DOI:** 10.1101/2025.10.05.680589

**Authors:** Ahmad H. Kabir, Asha Thapa, Bishrant Pant, Maruf Khan, Shifat Ara Saiful, Shyamal K. Talukder

## Abstract

**Aims:** *Trichoderma afroharzianum* T22 is widely recognized for enhancing plant stress resilience, yet its effects in pea plants may vary depending on iron (Fe) availability.

**Methods and Results:** We assessed the impact of T22 on pea grown under differential Fe status through integrated physiological and omics analyses. We found that the benefits of T22 are highly context dependent, providing significant improvements in photosynthesis and Fe/N accumulation under Fe deficiency but minimal effects under sufficiency. RNA-seq identified 262 DEGs under Fe deficiency and 555 DEGs under Fe sufficiency following T22 inoculation, with the latter primarily associated with basal metabolic functions, indicating potential colonization costs rather than adaptive responses. Particularly, T22 inoculation upregulated symbiosis-related genes (*Nodule-specific GRPs*, *Major facilitator, sugar transporter-like*), Fe transporters (*NRAMPs*, *HMAs*), and redox-associated genes (*Glutathione S-transferase*, *Glutathione peroxidase*) in the roots under Fe shortage, reflecting a coordinated response to enhance nutrient acquisition and stress tolerance. Microbiome profiling revealed that under Fe deficiency, T22 reshaped the root community by enriching several bacterial taxa such as Comamonadaceae, Pseudomonadaceae and *Mitsuaria*. These enriched bacterial taxa may act as potential ‘helpers’ to T22 by providing complementary thereby amplifying its beneficial effects under Fe deficiency. In contrast, under Fe sufficiency, community restructuring was primarily limited to the enrichment of Rhizobiaceae, *Pararhizobium*. Fungal taxa showed minimal overall changes, with the exception of a significant enrichment of *Paecilomyces* in response to T22 under Fe-deficient soil conditions.

**Conclusions:** These findings indicate that T22 functions in a context-dependent manner, with bacterial enrichment varying with Fe availability, while fungal helper effects were not prominent following T22 inoculation in pea plants.

**Impact Statement:** Beneficial microbes do not function uniformly across environments. This study highlights how Fe availability determines whether *Trichoderma* acts as a strong mutualist or a neutral colonizer in pea plants. Our findings advance a precision bioinoculant framework for deploying microbial consortia specifically in Fe-deficient soils to enhance legume resilience and sustainable crop production.

## 1. Introduction

Iron (Fe) deficiency is a pervasive constraint in legume crops grown on calcareous and alkaline soils, where high pH and carbonate content reduce Fe solubility and drive leaf chlorosis and yield losses (Merry et al., 2022; Assefa et al., 2020). Fe is indispensable for photosynthesis, respiration, and redox homeostasis, and therefore for biomass production and crop quality (Briat 2015). In legumes, Fe nutrition carries additional significance because nodulation and symbiotic N fixation require abundant Fe for nitrogenase, leghemoglobin, and other Fe cofactors (Liu et al., 2023; Brear et al., 2013). Consequently, Fe deficiency impairs nodule initiation and function and reduces whole-plant nitrogen status (Brear et al., 2013).

To acquire scarcely soluble Fe, dicots including legume deploy the Strategy I reduction mechanism acidifying the rhizosphere, reducing Fe(III) to Fe(II), and importing Fe(II), while grasses use Strategy II chelation with phytosiderophores; convergence and crosstalk between these strategies are increasingly recognized (Jeong and Guerinot 2009; Staiger, 2002). However, due to the lack of popular high-yield varieties with tolerance strategies, it remains a persistent issue for legume and other crops. Fe-deficiency-induced chlorosis management in crops typically relies on genotype selection and soil/seed treatments (Liberal et al., 2020). Yet, durable, biologically based solutions that enhance Fe bioavailability and maintain root–nodule performance remain a priority.

Harnessing the plant-associated microbiome could offer a promising alternative to conventional approaches (Compant et al., 2025). *Trichoderma* spp. are ubiquitous root-associated fungi widely used as biologicals that stimulate plant growth and resilience and changes to the rhizosphere microbiome (Thapa et al., 2025; Geng et al., 2025). Several *Trichoderma* strains produce and/or stimulate siderophores and organic acids that can increase Fe solubility for alleviating chlorosis induced by Fe deficiency (Thapa et al., 2025; Kabir and Bennetzen 2024; Tyśkiewicz et al., 2022). In general, plant–microbe mutualisms are context dependent; the magnitude and nature of the benefits vary with environmental conditions, host genotype, and microbial community composition (Thapa et al., 2025; Brown et al., 2020). However, these interactions are not static; under nutrient-rich conditions, the cost of maintaining the symbiont may outweigh its benefits, shifting the relationship away from mutualism (Kiers et al., 2003). Environmental stresses can further modulate mutualistic outcomes, often enhancing the reliance on microbial partners for nutrient acquisition and stress mitigation (Lamers et al., 2020; de Zelicourt et al., 2013). Understanding this context dependency is crucial for designing microbial inoculants that deliver consistent benefits under variable field conditions.

Despite the promise of *Trichoderma*, it remains unclear how Fe availability modulates the behavior of *Trichoderma afroharzianum* T22. Here we explicitly ask how T22 behaves differently in the presence versus absence of Fe deficiency at the root-soil interface. Using peas (*Pisum sativum*) as the host, we studied the differential effect of T22 on physiological responses paired with host transcriptomics and microbial community profiles in the rhizosphere of pea plants under Fe-sufficient and Fe-deficient conditions. This approach provides a systems view of how a well-characterized beneficial fungus recalibrates pea performance across Fe regimes.

## 2. Materials and methods

### 2.1. Plant cultivation and growth conditions

Garden pea (Sugar Snap variety) seeds were surface disinfected by soaking in a 1% sodium hypochlorite solution for 5 min, followed by thorough rinsing with sterile distilled water. Germination was performed in trays placed in an incubator at 25 °C for 2d Seedlings were transplanted into pots (500g soil/pot) containing a 1:3 mixture of field soil and SunGro potting mix and grown in the growth chamber in two growth conditions (Fe-sufficient and Fe-deficient supplemented with 4.5 g CaCO_3_ and 7.5 g NaHCO_3_ per pot) with or without the inoculation of *Trichoderma afroharzianum* T22 (1 × 10^9^ cfu/gram). The field soil was collected from the Scott Research and Extension Center in Winnsboro, Louisiana and characterized at the Agricultural & Environmental Services Laboratories, University of Georgia (Supplementary Table S1). The experiment was conducted following a randomized complete block design (RCBD) with three individual biological replicates (individual plant) per treatment for up to 5 weeks after transplantation. Plants were grown in a controlled growth chamber under a 10 h light/14 h dark photoperiod, with a light intensity of 250 μmol m ² s ¹ at ∼25 °C.

### 2.2. Determination of morpho-physiological attributes

The plant height was measured from below ground root to the highest leaf using a measuring tape immediately before harvesting the plants. The aboveground portions of the plant were harvested, and fresh biomass weight was measured using a scale. Nodules were counted manually on the roots of pea plants after carefully washing off adhering soil under running water. SPAD score, an indicator of chlorophyll content in fully expanded leaves, was measured using a handheld SPAD meter (AMTAST, United States). Also, Fv/Fm (maximal photochemical efficiency of PSII) was measured on the uppermost fully expanded leaves after dark adaptation for 1 hour, using a FluorPen FP 110 (Photon Systems Instruments, Czech Republic).

### 2.3. Fe and N analysis of plant tissues

After harvest, the roots of pea plants were carefully excised and rinsed under running tap water to remove adhering soil. The roots were then immersed in 0.1 mM CaSO for 10 min to displace surface ions, followed by a final rinse with deionized water. Roots and leaves were oven-dried at 75 °C for three days, and the dried samples were submitted to the Agricultural & Environmental Services Laboratories, University of Georgia, for iron quantification using inductively coupled plasma mass spectrometry (ICP-MS). Fe and N concentrations were determined by comparing ion signal intensities with those of certified calibration standards.

### 2.4. Molecular detection of T22 colonization in the roots

Colonization of T22 in pea root tips was assessed by quantifying the relative abundance of the T22 marker gene *TaAOX1* (L-amino acid oxidase) using DNA-based qPCR (Kabir and Bennetzen, 2024). Root tips (∼0.2 g) were washed twice in sterile phosphate-buffered saline, vortexed briefly, rinsed twice with sterile water, and subjected to DNA extraction using the CTAB method (Clarke, 2009). DNA was quantified (NanoDrop ND-1000, Wilmington, USA) and normalized across samples. qPCR was performed on a CFX96 Touch Real-Time PCR System (Bio-Rad, USA) with *TaAOX1*-specific primers (forward: 5′-GTC GGT AGC TGA AAG GGG AT-3′; reverse: 5′-ATT AGG CCG GAA ACA CC-3′). All reactions were run with iTaq Universal SYBR Green Supermix (Bio-Rad, USA), and *PsGAPDH* was used as the internal control for relative quantification by the 2−ΔΔCT method (Livak and Schmittgen, 2001). Relative abundance of *TaAOX1* was calculated using the 2 ΔΔCT method, normalized to the internal reference gene *PsGAPDH*, and expressed relative to the non-inoculated control root samples.

### 2.5. Determination of siderophore levels in the rhizosphere

Siderophore content in rhizosphere soil was quantified using the chrome azurol S (CAS) assay (Himpsl and Mobley, 2019). Briefly, soil samples were homogenized in 80% methanol and centrifuged at 10,000 rpm for 15 min. The supernatant (500 μL) was mixed with 500 μL CAS solution and incubated at room temperature for 5 min. Absorbance was recorded at 630 nm, with 1 mL CAS reagent serving as the reference. Siderophore units were calculated as %SU = [(Ar − As)/Ar] × 100, where Ar is the absorbance of the reference and As is the absorbance of the sample.

### 2.6. RNA-seq analysis in the roots

RNA-seq was performed in the root samples. Root samples were washed twice in sterile phosphate-buffered saline (PBS) by vortexing for 10 s, followed by two rinses with sterile water to remove surface contaminants. Samples were ground in liquid nitrogen with a pre-cooled mortar and pestle, and the resulting powder was used for RNA extraction with the SV Total RNA Isolation System (Promega, USA) according to the manufacturer’s protocol. RNA integrity (RIN > 8) was confirmed, and 1 µg of total RNA per sample was used for library preparation with the KAPA HiFi HotStart Library Amplification Kit (Kapa Biosystems, USA). Libraries were sequenced on an Illumina NextSeq SE 100 bp platform, yielding >80% high-quality reads (Phred > 33 per sample).

Adaptor and low-quality sequences were trimmed using Trimmomatic v0.38 (Bolger et al., 2014). Filtered reads were aligned to the pea reference genome (Yang et al., 2022) with HISAT2 v2.0.5 (Kim et al., 2015). Gene-level read counts were obtained using HTSeq (Anders et al., 2014). Differentially expressed genes (DEGs) were identified with the Limma R package v3.19 (Law et al., 2016) following normalization with the FPKM method. DEGs were defined as genes with log2FC ≥ 1 or ≤ −1 and false discovery rate (FDR) ≤ 0.05. Heatmaps were generated using the *pheatmap* R package. Pea gene ontology annotations were retrieved from Ensembl Plants and enrichment analysis was performed with ShinyGO v0.80.

### 2.7. Amplicon sequencing of 16S and ITS microbial communities

Microbial communities in pea roots were analyzed by amplicon sequencing of the 16S rRNA gene (bacteria) and ITS region (fungi). Roots were cleaned by vortexing in sterile phosphate-buffered saline (PBS) for 10 s, followed by two rinses with sterile water to remove surface contaminants. DNA was extracted from ∼0.2 g of root tissue using the Wizard® Genomic DNA Purification Kit (Promega, USA), including RNase and Proteinase K treatments to remove RNA and protein impurities. The 16S rRNA gene library was prepared with primers 341F (CCT ACG GGNGGC WGC AG) and 805R (GAC TAC HVGGG TAT CTA ATC C), and sequencing was performed on an Illumina NovaSeq 6000 (PE250). Raw reads were quality trimmed with cutadapt (Martin, 2011) and processed using the DADA2 pipeline (Callahan et al., 2016). Sequences of mitochondrial and chloroplast origin were removed. Taxonomic classification of amplicon sequence variants (ASVs) was performed with the UNITE reference database (Nilsson et al., 2019) for ITS and Silva v138 (Quast et al., 2013) for 16S. Community structure was assessed by Principal Coordinate Analysis (PCoA) based on Bray–Curtis dissimilarity of Hellinger-transformed counts using the *phyloseq* and *vegan* R packages. Taxa absolute abundance differences were tested with the non-parametric Kruskal–Wallis test. Alpha diversity, beta diversity and relative abundance were conducted using *phyloseq* (McMurdie and Holmes, 2013), with significance determined at *p* < 0.05.

### 2.8. Statistical analysis

All measurements were conducted with a minimum of three independent biological replicates per treatment. Statistical significance (*p* < 0.05) was determined using Student’s *t*-test in Microsoft Excel, and data visualization was performed with the ggplot2 package in R. Furthermore, a visual illustration was prepared using BioRender.

## 3. Results

### 3.1. Growth, photosynthesis, and nutrient status

Under Fe sufficiency, T22 had no significant effect on any measured trait (Fig. 1A–H, ns throughout). Plant height and biomass trended slightly upward with T22 (Fig. 1A–B, ns), and leaf SPAD and Fv/Fm values were comparable between −T22 and +T22 (Fig. 1C–D, ns). Root and leaf Fe concentrations were likewise unchanged (Fig. 1E–F, ns), and tissue N in roots and leaves remained statistically indistinguishable (Fig. 1G–H, ns). In contrast, under Fe deficiency, T22 conferred significant improvements across scales. T22-inoculated plants were taller and accumulated more biomass than uninoculated controls (Fig. 1A–B, significant). Photosynthetic performance recovered with T22: SPAD values increased and Fv/Fm rose relative to −T22 (Fig. 1C–D, significant). T22 also elevated Fe status, raising Fe concentrations in both roots and leaves (Fig. 1E–F, significant). Finally, T22 improved plant nitrogen status under Fe stress, increasing N% in roots and leaves compared with −T22 (Fig. 1G–H, significant). Together, these patterns show that T22’s benefits are Fe-contingent—physiology and nutrient status are stabilized specifically under Fe limitation, while remaining non-significantly changed when Fe is sufficient.

**Fig. 1.**
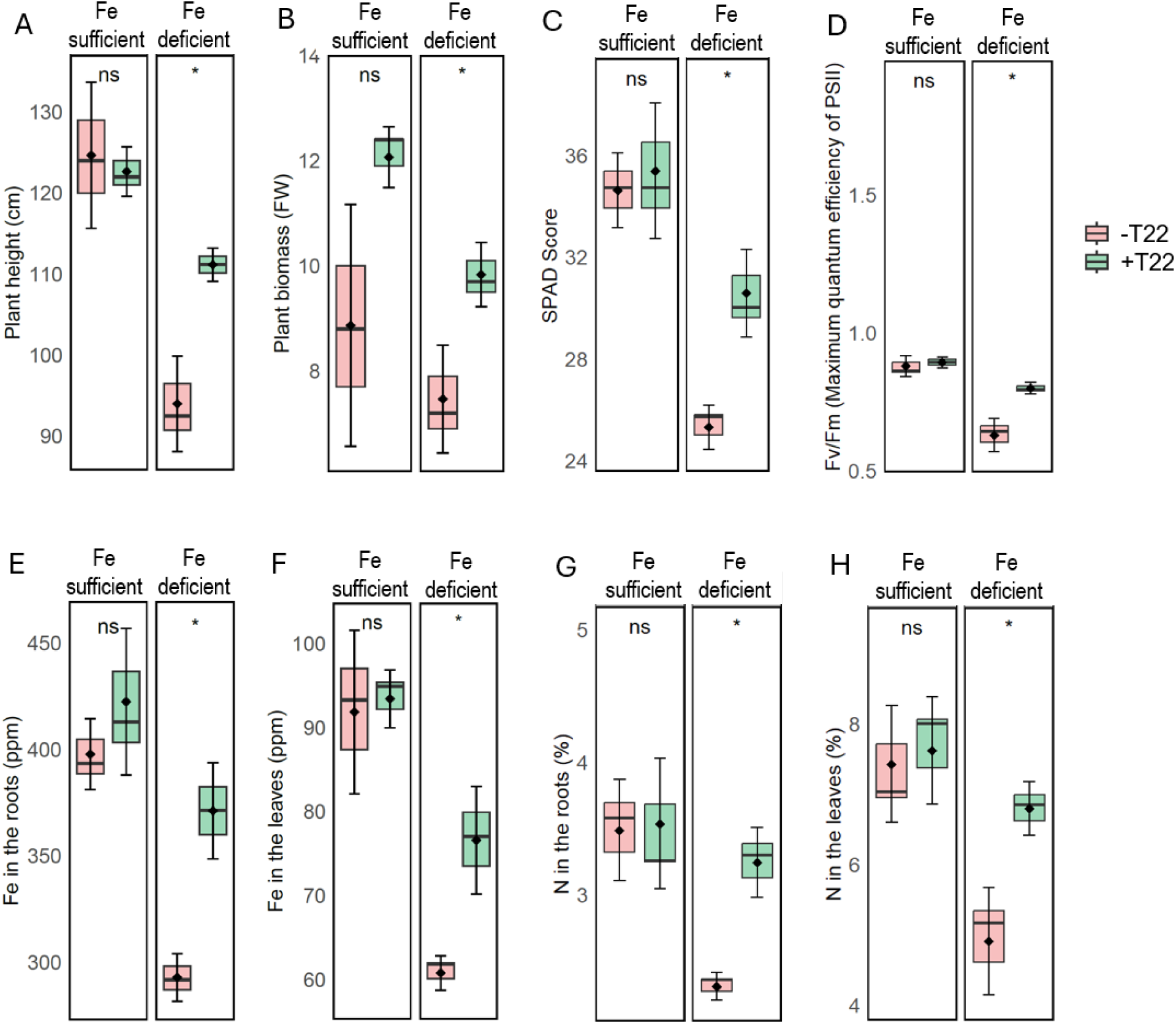
Effect of *Trichoderma afroharzianum* T22 under Fe-sufficient and Fe-deficient conditions on (A) Plant height, (B) plant biomass (fresh weight), (C) SPAD chlorophyll index, (D) maximum quantum efficiency of PSII (Fv/Fm), (E) Fe concentration in roots, (F) Fe concentration in leaves, (G) N concentration in roots, and (H) N concentration in leaves without (–T22) or with (+T22) inoculation in Sugar Snap. Boxplots show median (line), interquartile range (box), and range (whiskers); diamonds denote the mean. Pea plants were cultivated for 5 weeks; n = 3 independent biological replicates per treatment. Statistical differences between – T22 and +T22 within each Fe regime were assessed by two-tailed Student’s *t*-test (*p* < 0.05 indicated by “*”; ns, not significant).

### 3.2. T22 abundance, nodulation, and siderophore activity

Under Fe sufficiency, T22 inoculation significantly increased the relative abundance of T22 in the rhizosphere (Fig. 2A), while effects on nodules per plant and siderophore production were not significant (Fig. 2B–C). In contrast, under Fe deficiency, all three responses improved with T22: T22 abundance increased significantly (Fig. 2A), nodulation rose significantly (Fig. 2B), and siderophore production was significantly higher than −T22 (Fig. 2C).

**Fig. 2.**
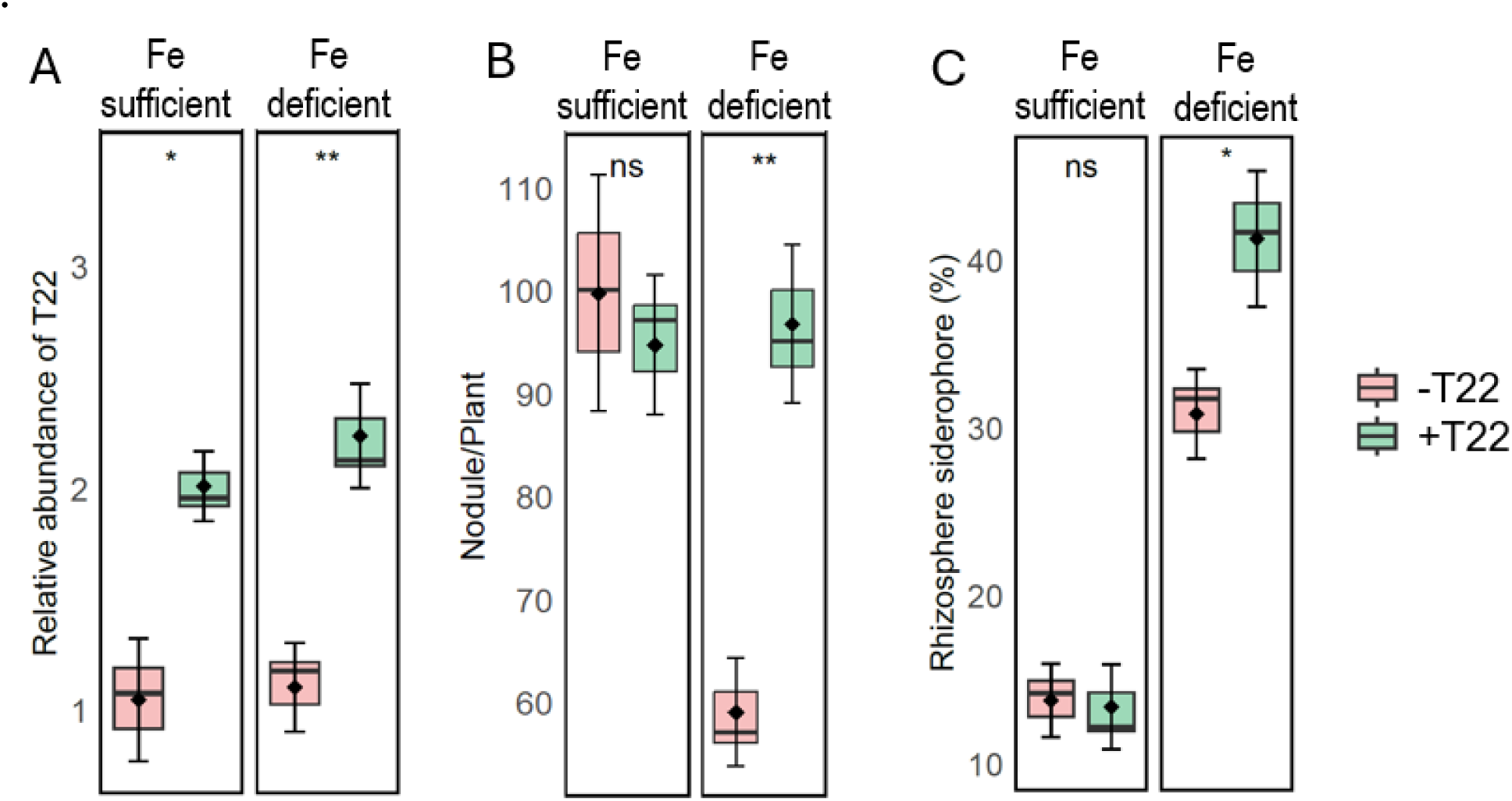
Effect of *Trichoderma afroharzianum* T22 inoculation under Fe-sufficient and Fe- deficient conditions on (A) Relative abundance of T22 normalized to the reference gene (*PsGAPDH*), (B) number of nodules per plant, and (C) rhizosphere siderophore production (%) with (+T22) or without (–T22) inoculation in Sugar Snap. Data represent mean ± SD from three independent biological replicates. Statistical differences between –T22 and +T22 within each genotype and Fe treatment were determined using a two-tailed Student’s *t*-test (*p* < 0.05, \**p* < 0.01; ns, not significant).

### 3.3. Differential gene expression

Across conditions, T22 reshaped the pea root transcriptome in an Fe-dependent manner (Fig. 3A–C). Under Fe sufficiency, we detected 555 DEGs relative to –T22 (225 upregulated, 330 downregulated; FDR < 0.05). Under Fe deficiency, fewer genes responded (262 DEGs; 95 up, 167 down; FDR < 0.05). Although the DEG count was larger in Fe sufficiency, the direction and magnitude of key Fe- and stress-associated genes shifted most clearly under Fe deficiency:

**Fig. 3.**
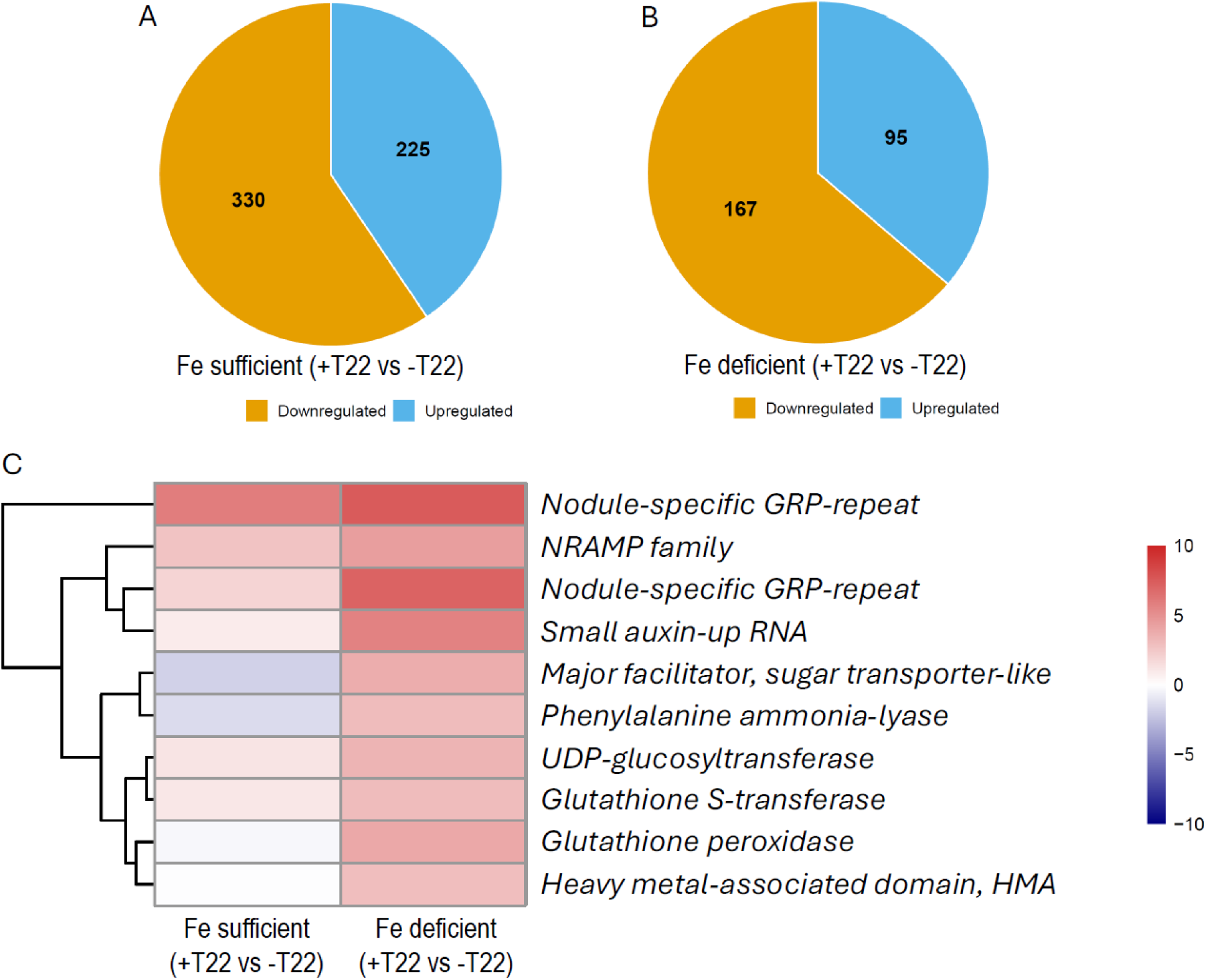
Transcriptomic changes induced by *Trichoderma afroharzianum* T22 inoculation under Fe-sufficient and Fe-deficient conditions. (A, B) Numbers of differentially expressed gene (DEGs) in roots of Fe-sufficient (A) and Fe-deficient (B) pea plants inoculated with T22 (+T22) compared to uninoculated controls (–T22) in Sugar Snap. Genes with |log fold change| ≥ 1 and adjusted *p* < 0.05 were considered significantly upregulated (blue) or downregulated (orange). (C) Heatmap of selected DEGs associated with nutrient transport, stress response, and secondary metabolism in Fe-sufficient and Fe-deficient conditions. Color scale represents log fold change values. Data were obtained from three independent biological replicates per treatment.

NRAMP transporters and HMA-domain metal-handling genes, a major facilitator/sugar-transporter–like gene, small auxin-up RNA (SAUR), phenylalanine ammonia-lyase (PAL), UDP-glucosyltransferase (UGT), and glutathione S-transferase (GST) all showed strong positive log FC with T22 specifically in Fe-deficient roots, whereas several of these were unchanged or reduced (not significant) under Fe sufficiency. Glutathione peroxidase displayed a weaker/negative shift under Fe sufficiency. Together, these patterns indicate that T22 preferentially activates Fe transport, hormone signaling, phenylpropanoid metabolism, and detoxification modules under Fe limitation, with many of the same loci not significantly induced when Fe is adequate (Fig. 3C).

### 3.4. GO enrichment highlights distinct functional pathways

GO over-representation recapitulated these contrasts (Fig. 4A–B). In Fe-sufficient plants, T22-responsive genes were significantly enriched for molecular functions related to nucleotide/ATP/adenyl-ribonucleotide binding, copper/anion/ion binding, transferase activity, and membrane terms (FDR < 0.05), while structural and transport categories were not significant. Under Fe deficiency, the program shifted: the top signals were cell-wall–related processes—structural constituent of cell wall (largest fold enrichment, >100), plant-type cell wall organization/biogenesis, and cell wall organization/biogenesis—accompanied by inorganic-ion transmembrane transporter activity, oxidoreductase activity, and metal/cation binding (all significant, FDR < 0.05). GO terms outside these sets were not significantly enriched. Overall, T22 elicits a condition-specific transcriptional response, emphasizing cell-wall remodeling and ion transport under Fe stress, but nucleotide-binding and general metabolic functions when Fe is sufficient (Fig. 4).

**Fig. 4.**
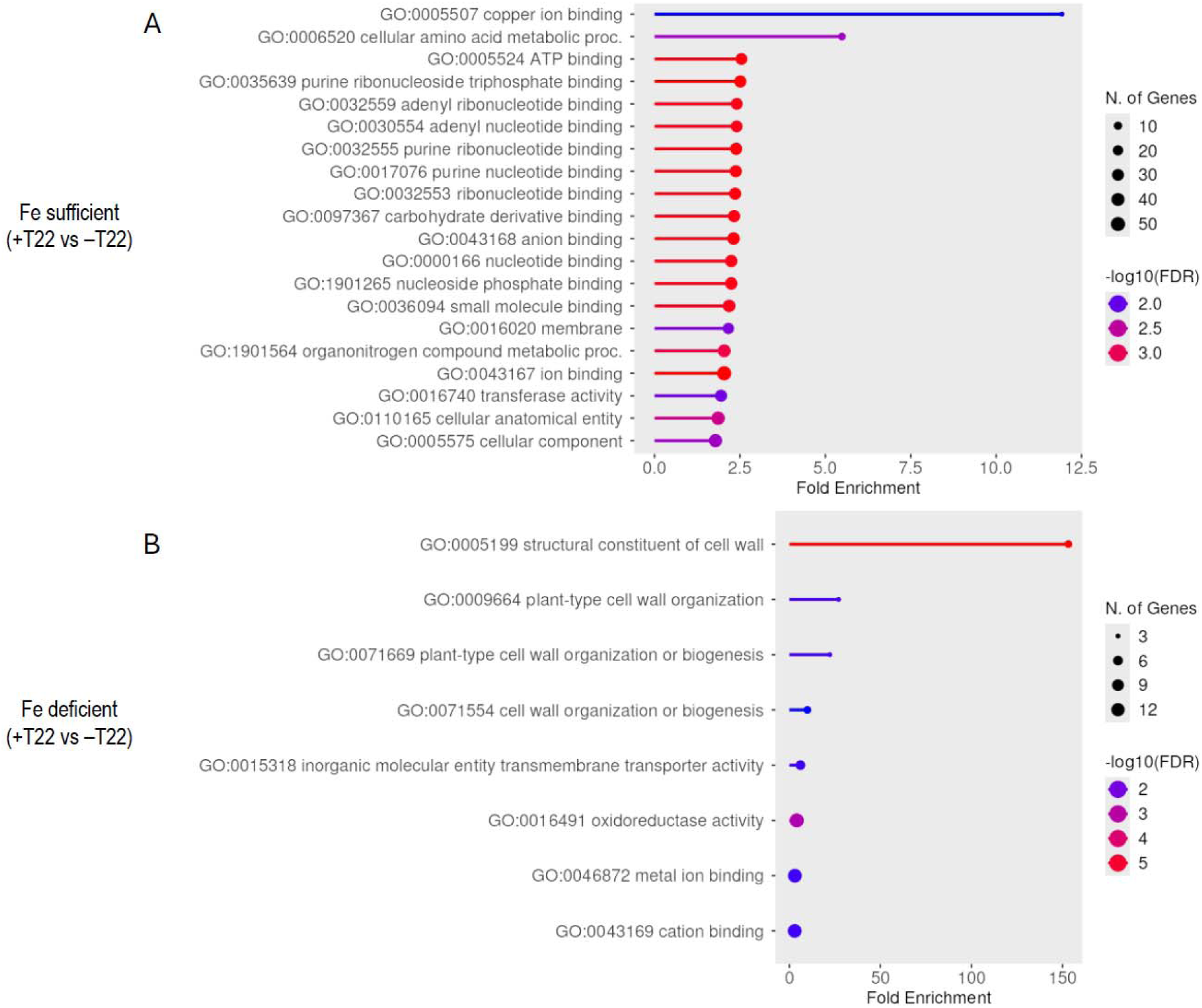
Gene ontology (GO) and enrichment analysis of differentially expressed genes in roots of plants inoculated with *Trichoderma afroharzianum* T22 (+T22) compared to uninoculated controls (–T22) under Fe-sufficient (A) and Fe-deficient (B) conditions in pea plants. The top enriched GO terms are shown for each condition, with fold enrichment on the x-axis. Dot size indicates the number of genes associated with each GO term, and dot color represents – log (FDR) values. Data was generated from three independent biological replicates per treatment.

### 3.5. Changes in bacterial community

Under Fe-sufficient conditions, alpha diversity (Chao1 richness and Simpson index) did not differ significantly between −T22 and +T22 treatments (*P* = 0.30 for both indices; Fig. 5A). Under Fe-deficient conditions, Chao1 richness remained non-significant between treatments (*P* = 0.32), whereas the Simpson diversity index showed a significant difference between −T22 and +T22 (*P* = 0.008; Fig. 5D). Under Fe-sufficient conditions, some taxa showed specific differences between −T22 and +T22 treatments. At the family level, Rhizobiaceae was higher in the +T22 treatment compared to −T22. At the genus level, *Pararhizobium* was also higher in +T22 relative to −T22. Under Fe-deficient conditions, only a subset of taxa showed significant differences between −T22 and +T22 treatments. At the family level, Comamonadaceae and Pseudomonadaceae were significantly increased in the +T22 treatment compared to −T22. At the genus level, *Mitsuaria* and *Variovorax* showed a significant increase and decrease, respectively, in +T22 relative to −T22 conditions (Fig. 5E).

**Fig. 5.**
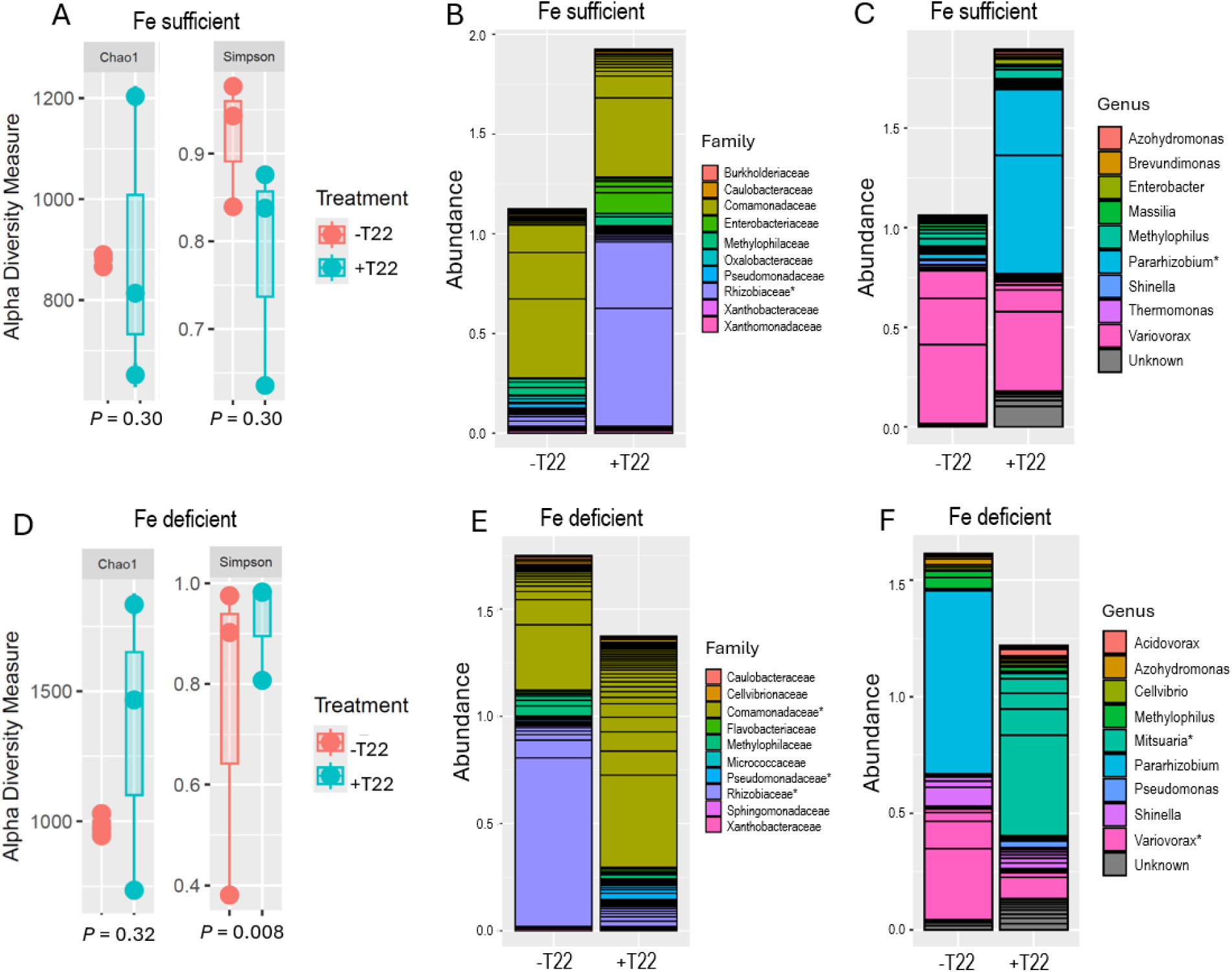
Effect of *Trichoderma afroharzianum* T22 inoculation on bacterial (16S) community diversity and composition in roots under Fe-sufficient and Fe-deficient conditions. (A, D) Alpha diversity indices (Chao1 richness and Simpson diversity). (B, E) Absolute abundance of dominant bacterial families. (C, F) Relative abundance of dominant bacterial genera. Taxa showing significant differences between +T22 and −T22 within each Fe condition are indicated with an asterisk (*, *p* < 0.05). Data represents the mean of three independent biological replicates per treatment.

### 3.6. Changes in fungal community

Under Fe-sufficient conditions, alpha diversity indices (Chao1 richness and Simpson diversity) did not differ significantly between −T22 and +T22 treatments (*P* = 0.43 and *P* = 0.39, respectively; Fig. 6A). Similarly, under Fe-deficient conditions, no significant differences were observed in Chao1 richness (*P* = 0.48) or Simpson diversity (*P* = 0.88) between treatments (Fig. 6D). Under Fe-sufficient conditions, only a few taxa showed significant differences between −T22 and +T22 treatments. At the family level, the abundance of Cladosporiaceae was lower in the +T22 treatment compared to −T22. At the genus level, the abundance of *Paecilomyces* was higher in +T22 relative to −T22. Under Fe-deficient conditions, only a limited number of taxa exhibited significant differences between −T22 and +T22 treatments. At the family level, Nectriaceae showed higher abundance in the +T22 treatment compared to −T22. At the genus level, *Fusarium* was significantly higher in +T22 relative to −T22 conditions.

**Fig. 6.**
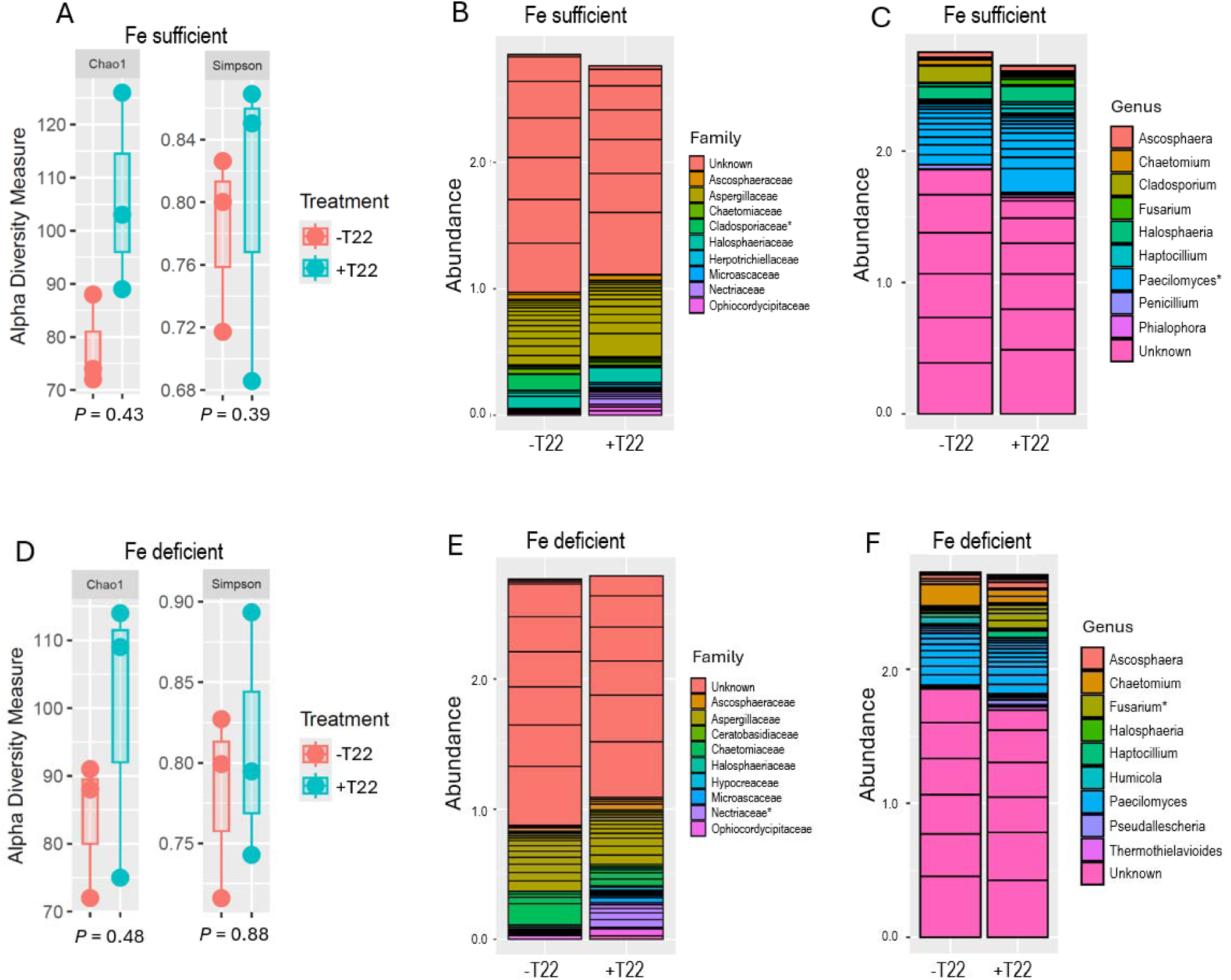
Effect of *Trichoderma afroharzianum* T22 inoculation on fungal (ITS) community diversity and composition in roots under Fe-sufficient and Fe-deficient conditions. (A, D) Alpha diversity indices (Chao1 richness and Simpson diversity). (B, E) Absolute abundance of dominant bacterial families. (C, F) Relative abundance of dominant bacterial genera. Taxa showing significant differences between +T22 and −T22 within each Fe condition are indicated with an asterisk (*, *p* < 0.05). Data represents the mean of three independent biological replicates per treatment.

## 4. Discussion

### 4.1. Context-dependent benefits of Trichoderma afroharzianum T22

Our study demonstrates that the benefits of T22 are not universal, but highly dependent on Fe nutritional status. Under Fe sufficiency, T22 colonized pea roots but did not significantly affect plant health parameters or Fe and N concentrations. By contrast, under Fe deficiency, T22 showed efficiency in improving these physiological and nutrient traits, including recovery of photosynthesis and Fe/N accumulation. This confirms the hypothesis that *Trichoderma*-plant interactions are conditional mutualisms, where benefits are realized primarily under nutrient stress (Donoso et al., 2008; Kiers et al., 2003). Similar context-dependent outcomes have been observed in other legumes. For instance, soybean grown in calcareous soils showed greater reliance on microbial partners under Fe stress, but minimal response when Fe was sufficient (Roriz et al., 2023; Assefa et al., 2020). These findings underscore that the symbiotic relationship is maintained by a cost–benefit balance, where plants minimize investment in microbial interactions when internal Fe status is adequate.

Although the functional effects of T22 were dependent on Fe availability, our data indicate that T22 colonization itself was not constrained by Fe status. This suggests that colonization is a context-independent trait, reflecting the strong colonization capacity of *Trichoderma* to penetrate and persist in root tissues regardless of nutrient status (Compant et al., 2025; Tyśkiewicz et al., 2022). This separation between colonization capacity and functional outcomes emphasizes that while *Trichoderma* can establish robustly in diverse conditions, the ecological and physiological benefits to the host depend on stress context. Similar findings have been reported in sorghum and peas, where T22 colonized roots broadly but only conferred growth or nutrient benefits under stress conditions (Kabir & Bennetzen, 2024; Thapa et al., 2025).

### 4.2. Fe deficiency unlocks the functional potential of T22

While T22 colonization occurred regardless of Fe availability, its functional consequences were strongly amplified in Fe-limited plants. Specifically, T22 colonization, siderophore release, and nodulation all increased significantly under Fe deficiency but remained unchanged under sufficiency. This suggests that the root–soil environment under Fe limitation actively promotes microbial functions that mobilize Fe. *Trichoderma* is well documented to produce siderophores and organic acids, which solubilize otherwise insoluble Fe in alkaline soils (Kabir and Bennetzen, 2024; Sood et al., 2020). Kabir and Bennetzen (2024) demonstrated that T22 enhances siderophore activity in sorghum, directly increasing Fe uptake and conferring Fe- deficiency tolerance. Our pea data supports this mechanism, as T22-inoculated plants displayed elevated siderophore activity and restored Fe concentrations in roots and leaves. Fe is indispensable for rhizobial symbiosis, since both nitrogenase and leghemoglobin require high Fe inputs (Lindström and Mousav, 2020; Brear et al., 2013). By mobilizing Fe, T22 may indirectly sustain nodulation, which in turn improves N acquisition supported by higher N concentrations in roots and shoots. This interaction represents a tripartite synergy among *Trichoderma*, *Rhizobium*, and the host, consistent with recent findings that T22 induces flavonoid-driven nodulation signals in pea under alkaline stress (Thapa et al., 2025). Moreover, studies suggest that plant growth and transcriptional responses to T22 are strongly influenced by substrate type and plant genotype (Thapa et al., 2025; Schmidt et al., 2020).

### 4.3. T22 Rewires root transcriptomes for Fe stress management

RNA-seq analysis revealed a distinct and Fe-dependent transcriptional reprogramming in pea roots colonized by *Trichoderma afroharzianum* T22. Under Fe deficiency, T22 strongly upregulated genes central to Fe uptake and transport, including *NRAMP* family members and heavy metal–associated (HMA) domain proteins. Similar transcriptional activation of NRAMPs has been reported in sorghum inoculated with T22, where enhanced Fe transport alleviated chlorosis symptoms (Kabir & Bennetzen, 2024). This activation aligns with the enhanced siderophore-mediated Fe mobilization observed in the rhizosphere, suggesting that T22 inoculation primes both fungal- and plant-mediated Fe acquisition strategies (Tyśkiewicz et al., 2022; Geng et al., 2025).

In addition, genes associated with phenylpropanoid metabolism (e.g., *phenylalanine ammonialyase*, PAL), hormone signaling (*small auxin-up RNA*, SAUR), and antioxidant defenses (*glutathione S-transferase*, GST; *glutathione peroxidase*, GPX) were activated. Phenylpropanoid and flavonoid pathways are widely implicated in signaling and microbial recruitment in legume roots (Thapa et al., 2025), while PAL-mediated phenylpropanoid flux has been shown to enhance both defense and symbiotic signaling under abiotic stress (Yadav et al., 2020). Similarly, GSTs and GPXs are well-established mediators of redox homeostasis in plants colonized by beneficial fungi, buffering oxidative stress generated during root colonization (Casatejada et al., 2023). The combined activation of these pathways suggests that T22 integrates nutrient uptake with redox regulation and stress signaling. Gene ontology enrichment further confirmed the preferential induction of pathways associated with cell-wall remodeling, oxidoreductase activity, and ion transport specifically under Fe limitation (Fig. 4B). These processes are consistent with reports that root cell-wall dynamics and oxidoreductase activity are central components of Strategy I Fe uptake responses in dicots (Hantzis et al., 2018; Jeong & Guerinot, 2009). Similarly, the induction of phenylpropanoid pathways and flavonoid metabolism has been widely linked to rhizosphere signaling and the recruitment of beneficial microbiota (Thapa et al., 2025). Upregulation of PAL and sugar transporter-like genes in our dataset suggests that T22 may promote enhanced root exudation, a mechanism that fosters microbial enrichment and nutrient exchange in the rhizosphere (Lei et al., 2025). Thus, T22 appears to modulate both direct nutrient transport and indirect exudate-mediated interactions to support Fe stress resilience.

In contrast, under Fe sufficiency, T22 inoculation triggered a greater number of DEGs (555 vs. 262 under deficiency), but these were largely associated with basal metabolic processes, such as nucleotide binding, protein turnover, and ion binding (Fig. 4A). These transcriptional changes may reflect the metabolic costs of colonization rather than an adaptive, stress-specific response. The lack of enrichment in Fe transport or redox-related pathways suggests that the host perceives T22 as an additional metabolic load rather than a symbiotic partner when Fe is abundant. This contrast highlights a hallmark of conditional mutualism: plants preferentially invest in transcriptional and metabolic shifts to support microbial partners only when external nutrient availability is low (Blonde et al., 2025; Jones et al., 2019). While the present study provides a comprehensive analysis of host transcriptomic responses, it does not directly assess the transcriptional responses of T22 under varying Fe conditions. Therefore, it remains unclear whether T22 actively modulates other metabolic pathways. Future studies integrating dual RNA-seq may be encouraged to capture this bidirectional interaction and to better resolve the molecular mechanisms underlying the context-dependent behavior of T22. Although fungal gene expression was not directly measured, the observed increase in siderophore activity, improved Fe accumulation, and enrichment of siderophore-producing bacterial taxa collectively suggest that T22 participates in Fe-responsive functional network in the rhizosphere. Furthermore, our findings demonstrate that T22 functions as a molecular switch under Fe stress, activating targeted transcriptional programs that facilitate Fe uptake, redox homeostasis, and microbial recruitment.

### 4.4. Microbiome restructuring and selective enrichment of beneficial taxa

Microbiome profiling confirmed that *Trichoderma afroharzianum* T22 reshaped root-associated bacterial and fungal communities in a condition-specific manner. Under Fe-sufficient conditions, T22 treatment leads to a pronounced enrichment of Rhizobiaceae (family level) and *Pararhizobium* (genus level), suggesting enhanced recruitment of symbiotically and functionally important rhizosphere bacteria. Rhizobiaceae can act in improved nutrient acquisition and root system development (Masson-Boivin et al., 2009). At the genus level, the strong enrichment of *Pararhizobium* further supports this observation. *Pararhizobium* species are closely related to classical rhizobia and are known for their roles in nitrogen metabolism, and plant–microbe signaling (Debnath et al., 2023). Their enrichment under T22 treatment indicates that Trichoderma may facilitate a rhizosphere environment that favors beneficial symbiotic or associative interactions. Under Fe deficiency, pea plants inoculated with T22 appear to actively recruit bacterial taxa with established roles in Fe mobilization, hormone signaling, and plant growth promotion. In particular, T22 selectively enriched several families, such as Comamonadaceae and Pseudomonadaceae, as well as the genus *Mitsuaria*. The enrichment of Pseudomonadaceae and Comamonadaceae is particularly significant, as members of these families are widely recognized for their plant growth–promoting traits, including nutrient solubilization, phytohormone production, and biocontrol activity (Lugtenberg & Kamilova, 2009; Willems, 2014). For example, *Pseudomonas* species are widely reported to produce high-affinity siderophores such as pyoverdine, which mobilize Fe and improve Fe uptake under alkaline soils (Cornelis, 2010; Saha et al., 2016). Furthermore, *Mitsuaria* is increasingly associated with plant-beneficial consortia, particularly for phosphate solubilization and siderophore-mediated Fe acquisition (Zhang et al., 2019). The consistent enrichment of these taxa in Fe-starved, T22-inoculated plants strongly suggests that the fungus functions as a “microbial recruiter,” promoting a rhizosphere microbiome specialized for Fe mobilization and nutrient cycling. On the fungal side, T22 application induces noticeable shifts in the fungal community at the genus level, particularly affecting taxa such as *Paecilomyces* under Fe- sufficient conditions. *Paecilomyces* is recognized for promoting plant growth through mechanisms such as enzyme production, nutrient mobilization, and antagonistic activity against plant pathogens (Bilal et al., 2021). Interestingly, T22 increased richness only under Fe deficiency, with enrichment of *Fusarium* (Fig. 6F). While these genera may include opportunistic or weak pathogens, their enrichment under T22 inoculation likely reflects community restructuring that enhances microbial competition and antagonism, thereby limiting pathogen establishment (Abbas et al., 2022). Similar restructuring effects have been reported for *Trichoderma*-mediated rhizosphere modulation in cereals and legumes, where non-pathogenic fungal taxa occupy ecological niches to suppress disease outbreaks (López-Bucio et al., 2015; Harman et al., 2021). These results align with the broader concept that plant-associated microbiomes are not static but are actively recruited by hosts and their microbial partners in response to nutrient limitations (Berendsen et al., 2012; Trivedi et al., 2020). Taken together, our findings support a working model in which T22 may act as a microbiome modulator, indirectly enhancing Fe acquisition through recruitment of functionally complementary bacterial partners. While our microbiome analysis revealed enrichment of several bacterial genera, but their functional roles in this system remain putative and are inferred from known traits. Therefore, the present study establishes strong associative links between microbial enrichment and improved plant Fe status but does not experimentally validate causality. Future efforts may focus on synthetic community-based approaches, targeted inoculation, or microbial depletion assays to directly validate the roles of these taxa in Fe mobilization and plant stress tolerance.

## 5. Conclusion

This study demonstrates that the benefits of T22 in pea are not universal, but strongly dependent on Fe nutritional status. While T22 colonization was observed under both Fe sufficiency and deficiency, its functional outcomes diverged sharply. Under Fe sufficiency, T22 had minimal effects on plant physiology, nutrient status, or microbial restructuring, indicating that colonization alone is not sufficient to drive host benefit. By contrast, under Fe deficiency, T22 significantly improved growth-related traits, photosynthesis, and Fe and N accumulation, underscoring the conditional nature of *Trichoderma*–plant mutualisms. T22 inoculation promoted siderophore release and sustained nodulation, supporting a tripartite synergy among *Trichoderma*, *Rhizobium*, and the host. Transcriptomic profiling revealed the activation of Fe uptake, phenylpropanoid metabolism, and antioxidant defenses processes. Simultaneously, microbiome profiling demonstrated that T22 restructured the rhizosphere community, selectively enriching Comamonadaceae, Pseudomonadaceae and *Mitsuaria* (Fig. 7). Together, our findings provide clear evidence that T22 functions as a molecular and microbial switch, conferring pronounced benefits only when plants face Fe starvation. This work underscores the potential of *Trichoderma*-based inoculants as sustainable, microbiome-driven solutions to enhance crop resilience in nutrient-limited agroecosystems. Furthermore, these findings represent an important step toward a systems-level understanding of plant–fungus–microbiome interactions and may be further advanced by incorporating fungal omics approaches.

**Fig. 7.**
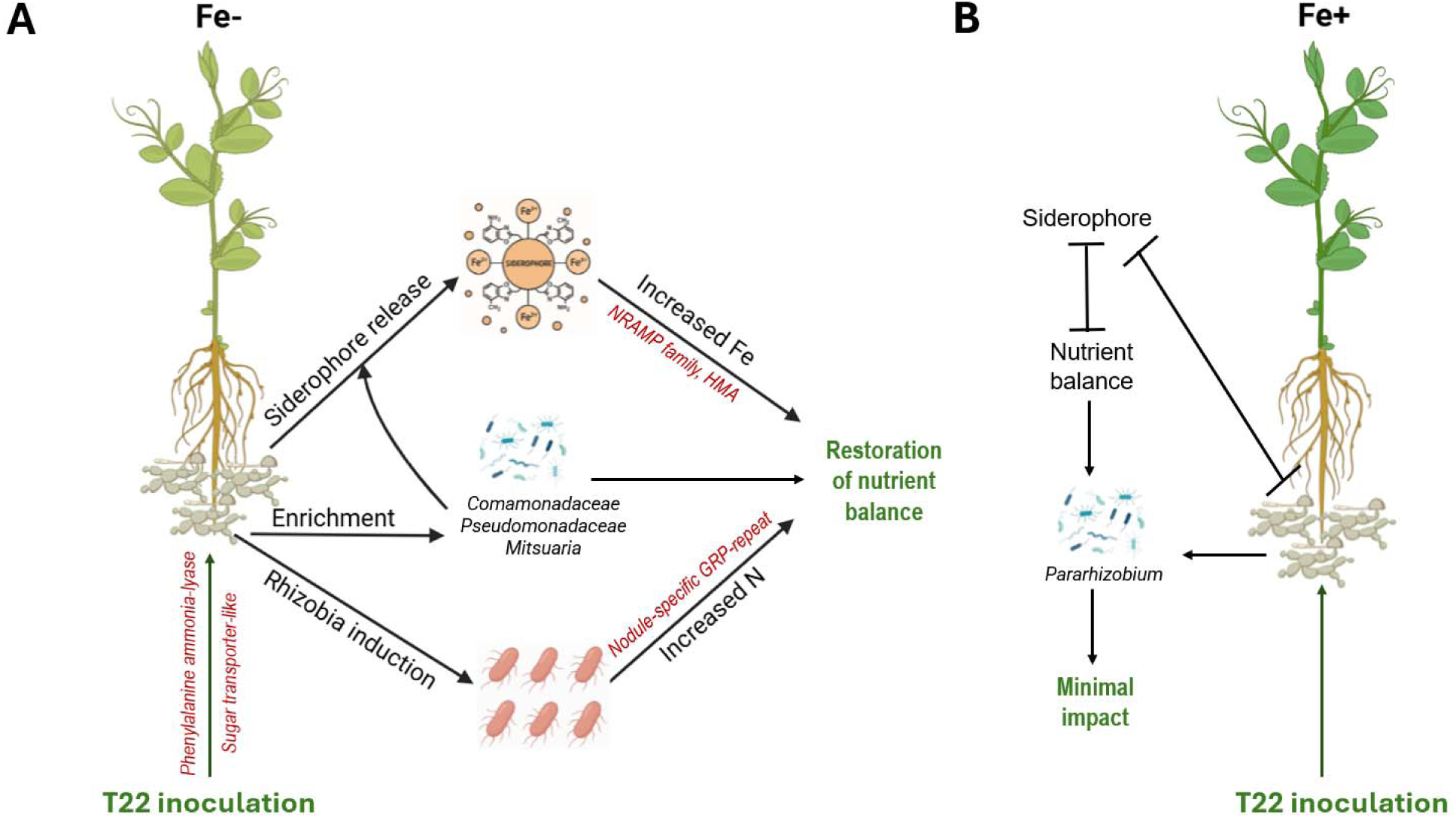
Model of *Trichoderma harzianum* T22–mediated effects on pea roots under Fe-deficient (Fe–) and Fe-sufficient (Fe+) conditions. (A) Under Fe deficiency, T22 inoculation leads to microbial enrichment of *Comamonadaceae, Pseudomonadaceae, Mitsuaria*. These change promote siderophore release and nodulation, resulting in increased Fe and N availability. Transcriptomic reprogramming further supports these processes, including the induction of *NRAMP* and heavy metal–associated (HMA) transporters for Fe uptake, nodule-specific GRP-repeat genes for N metabolism, phenylalanine ammonia-lyase for defense and metabolic adjustments, and sugar transporter-like genes for nutrient allocation. Together, these microbial and molecular responses restore nutrient balance under Fe limitation. (B) Under Fe sufficiency, T22 inoculation results in minimal microbiome restructuring, characterized by enrichment of *Pararhizobium*. Siderophore activity is repressed, transcriptional changes are weak, and overall contributions to Fe and N balance are limited, leading to minimal physiological impact.

## Author statement

We hereby certify that all authors have seen and approved the manuscript.

## Data availability

Illumina sequencing data were submitted to NCBI under the following accession numbers: PRJNA1313627 (RNA-seq), PRJNA1314923 (16S) and PRJNA1314929 (ITS).

## Ethical Approval Statement

This study did not involve human participants or vertebrate animals and therefore did not require approval from an institutional ethics committee.

## Acknowledgments

We express our gratitude to Louisiana Biomedical Research Network for the funding. This research was also supported by a startup grant from Lamar University.

## Author contributions

AHK conceived the study, performed the molecular experiments, analyzed the data, performed interpreted the results and drafted the manuscript. AT and MK helped with RNA-seq data analysis and revised the manuscript. SAS and BP assisted in amplicon data analysis and graphical representation. SKT interpreted the results and critically revised the manuscript.

